## Supplementary material for "Structural mechanisms for VMAT2 inhibition by tetrabenazine": Combined Figure Supplements

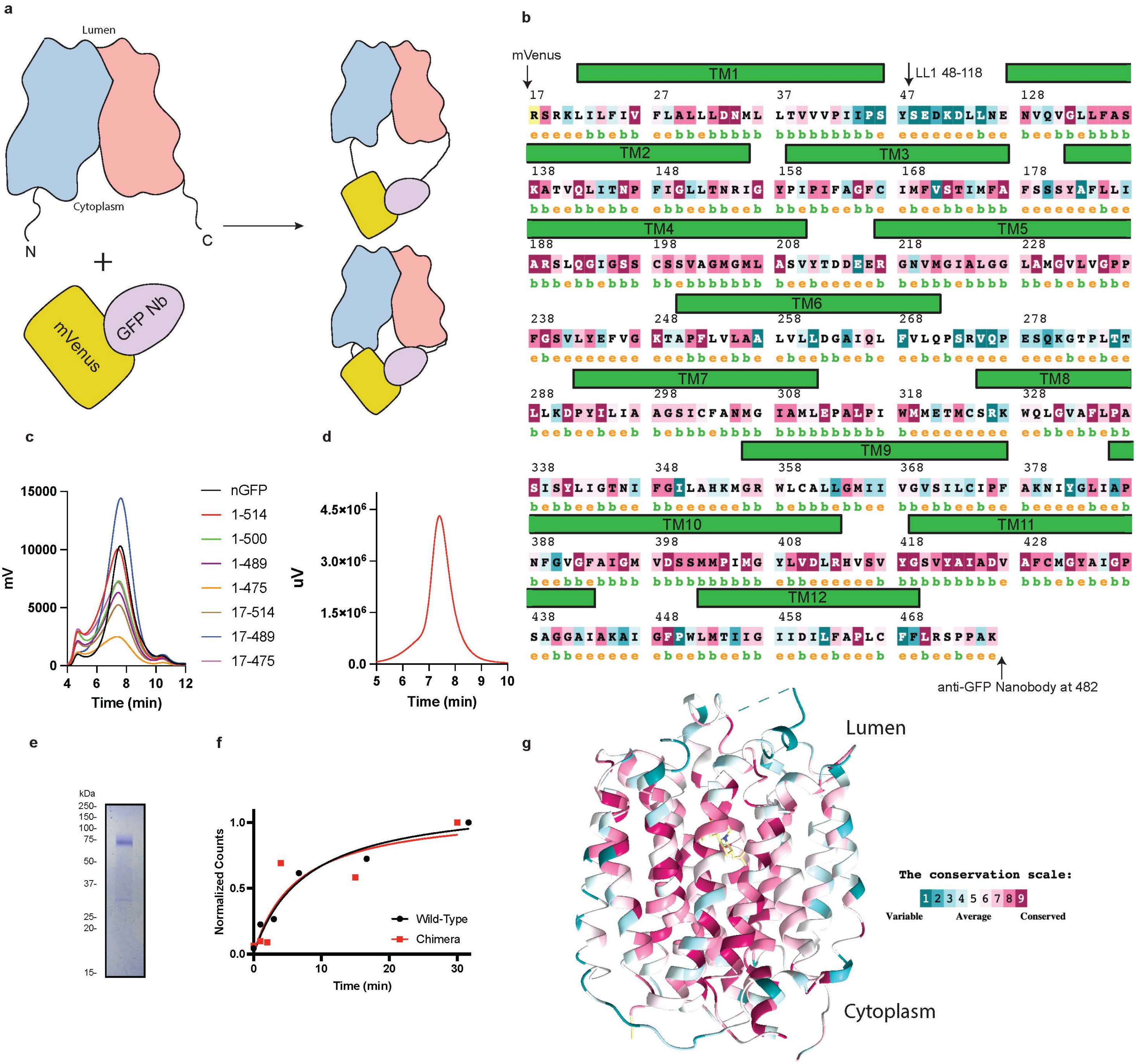


Supplementary Figure 1

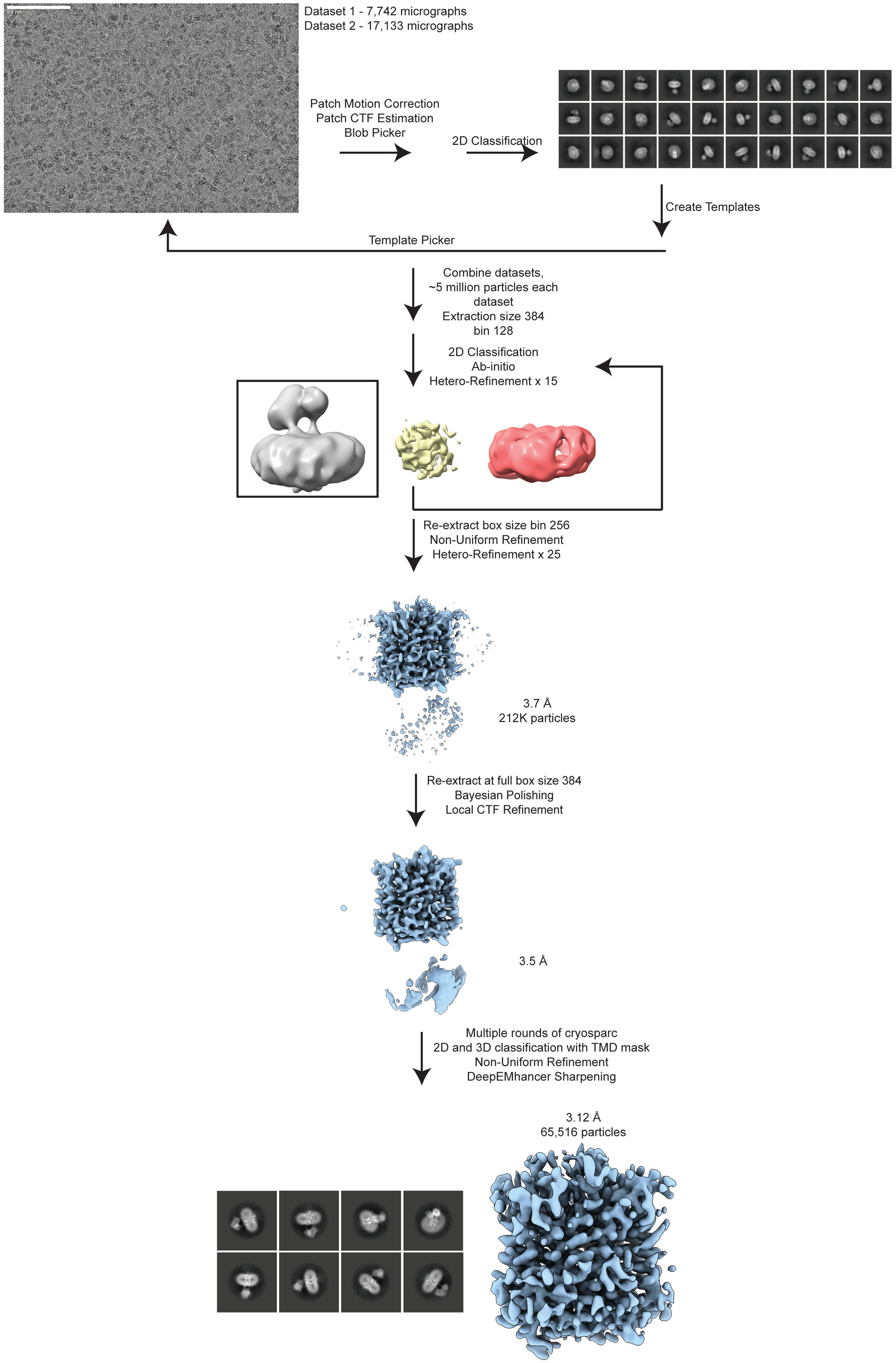


Supplementary Figure 2

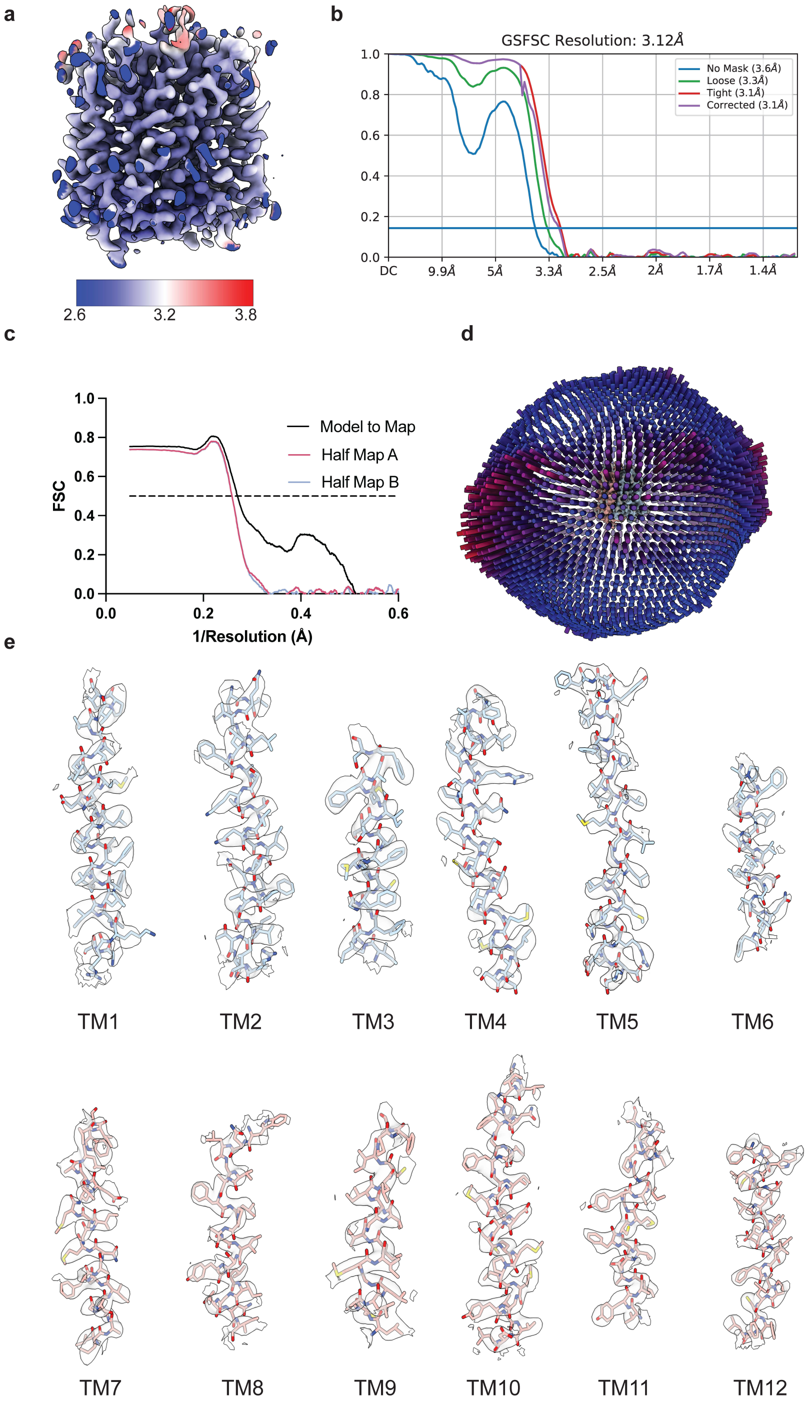


Supplementary Figure 3

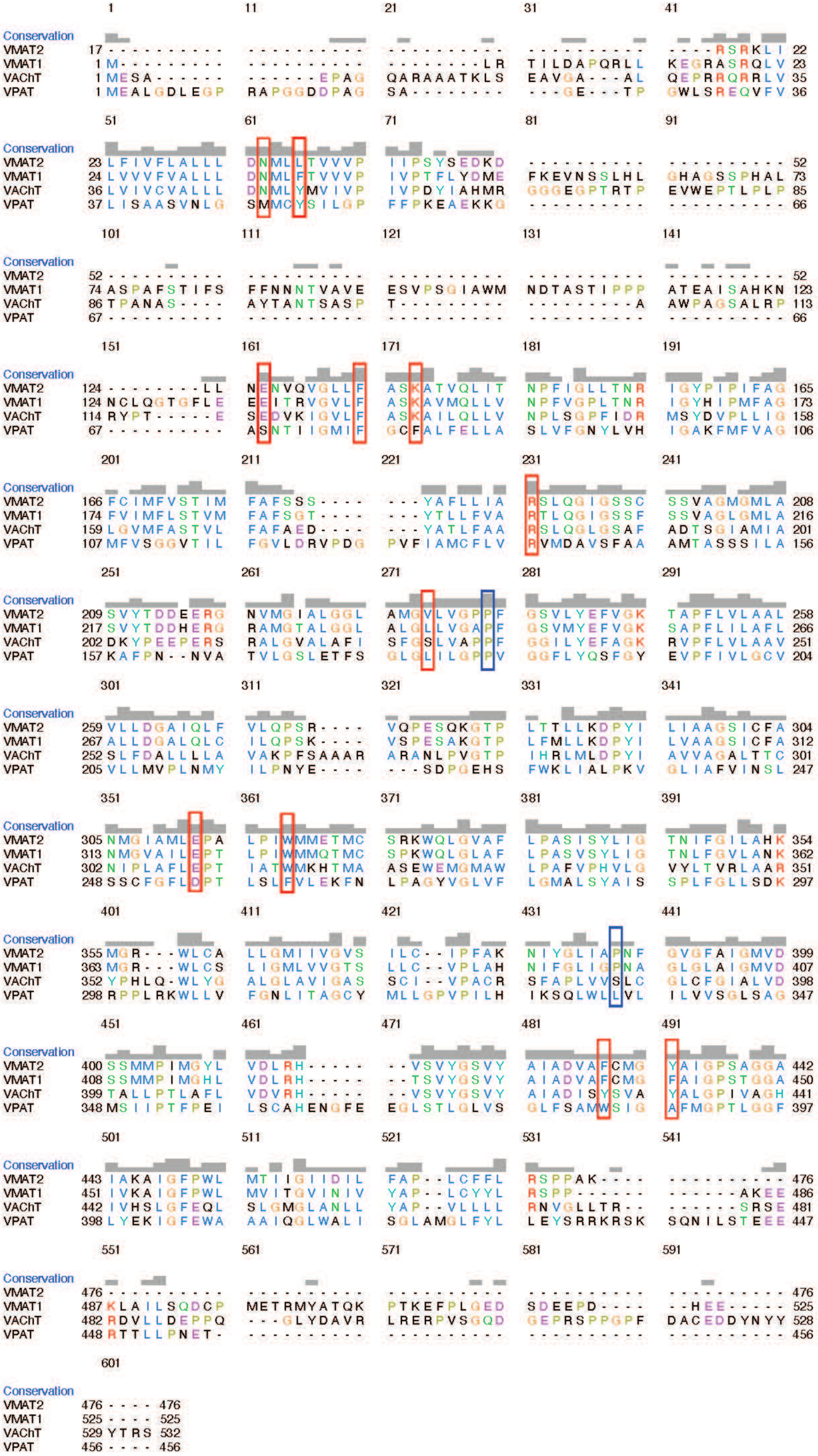


Supplementary Figure 4

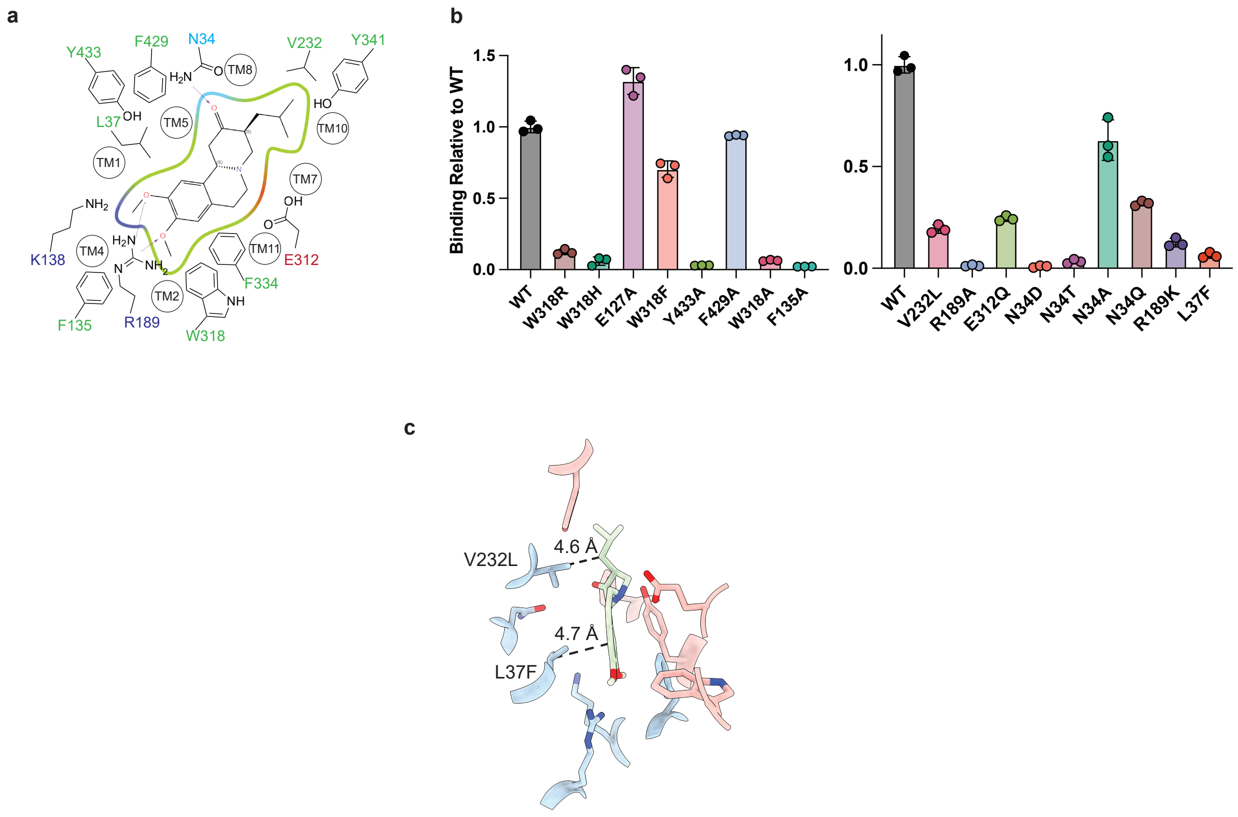


Supplementary Figure 5

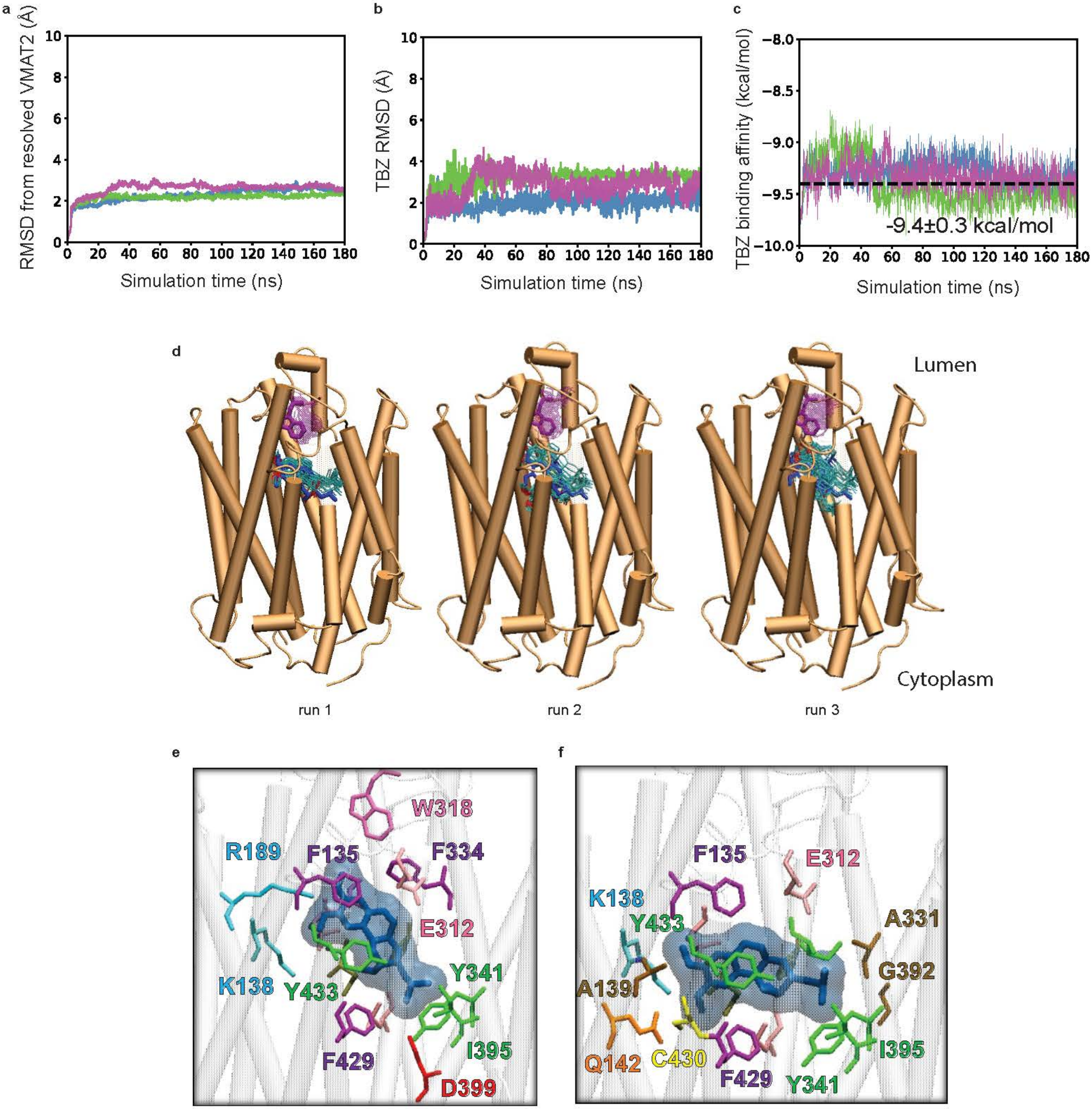


Supplementary Figure 6

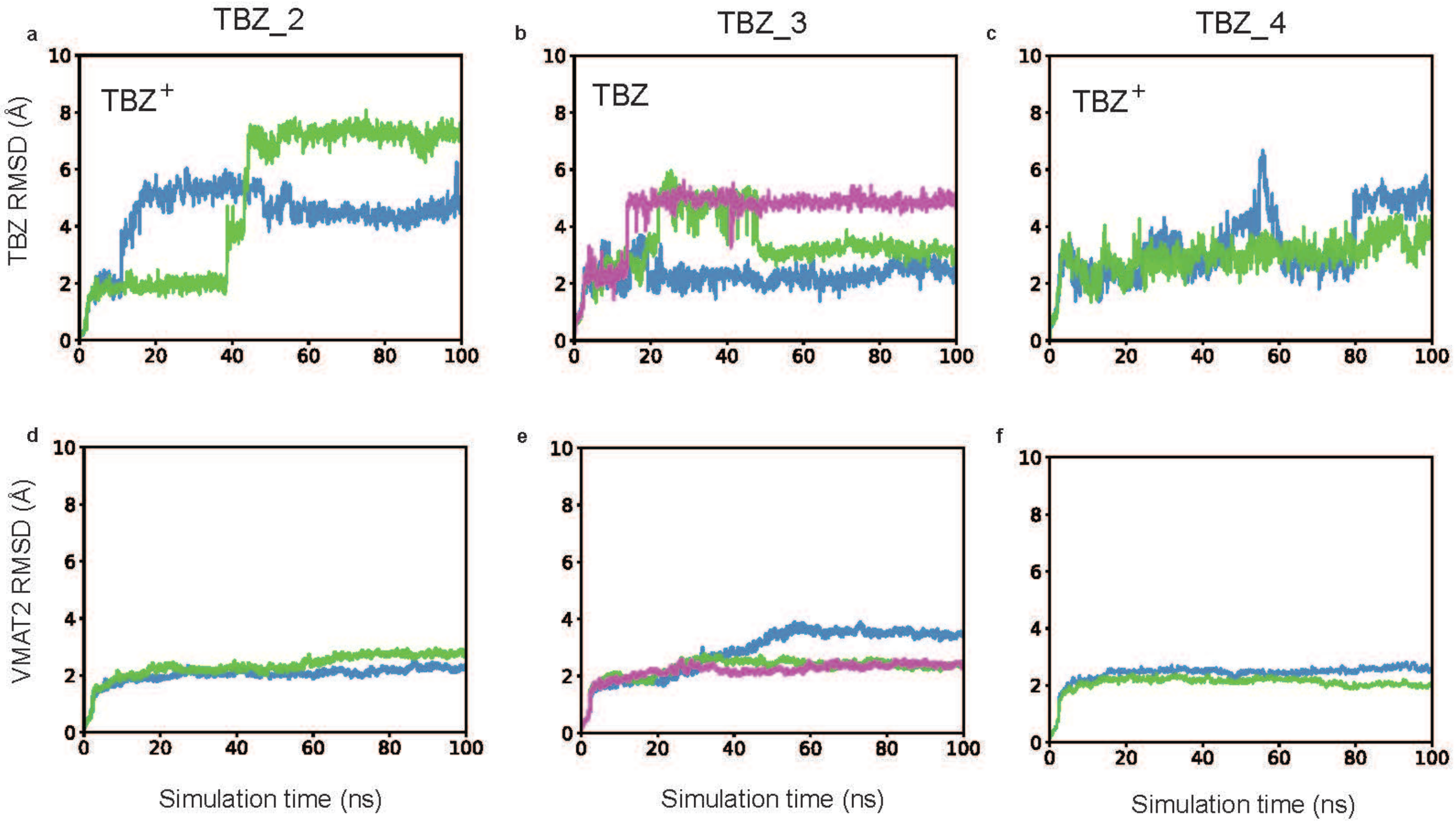


Supplementary Figure 7

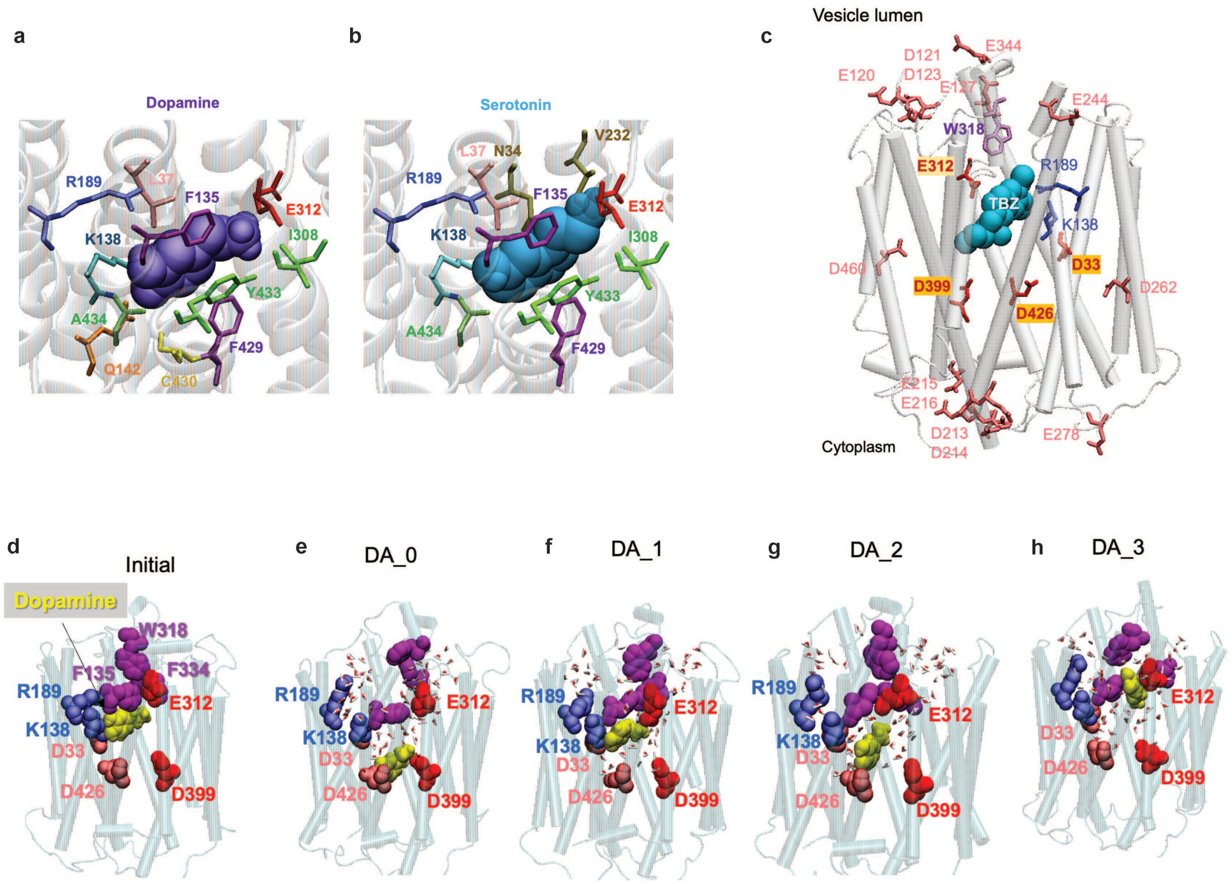


Supplementary Figure 8

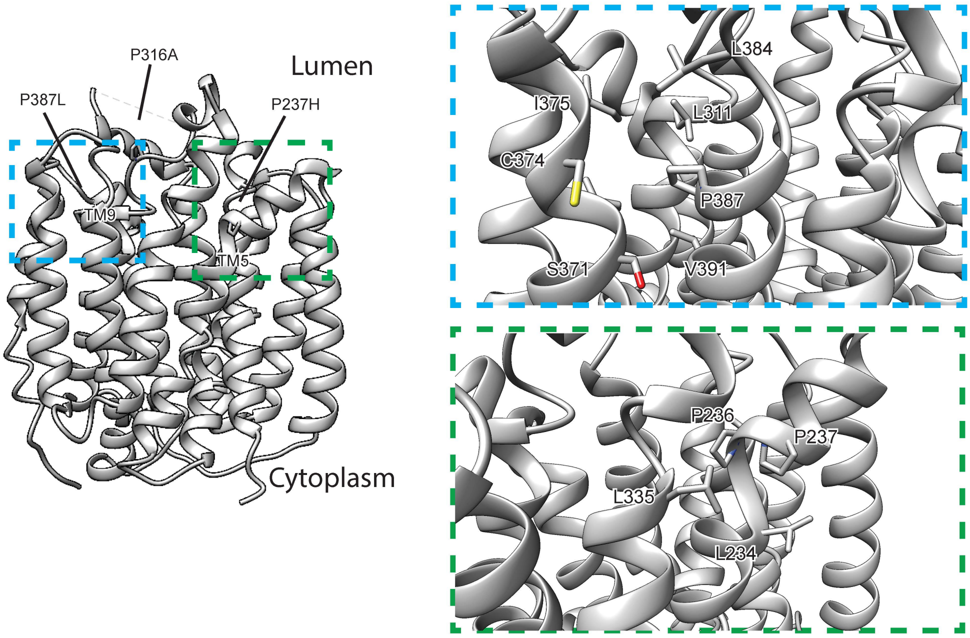


Supplementary Figure 9

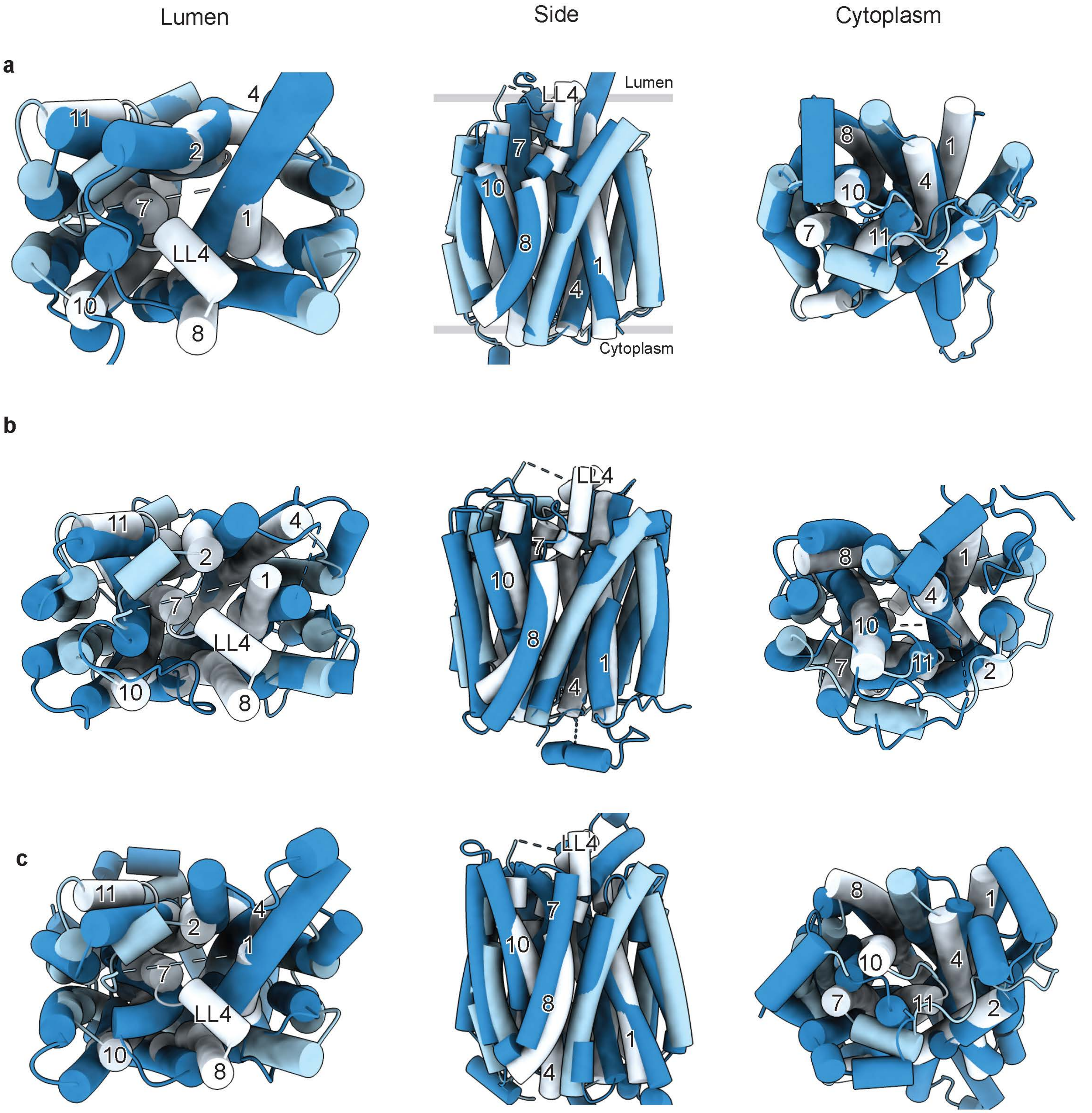


Supplementary Figure 10
